# Background selection from unlinked sites causes non-independent evolution of deleterious mutations

**DOI:** 10.1101/2022.01.11.475913

**Authors:** Joseph Matheson, Joanna Masel

**Affiliations:** Department of Ecology and Evolutionary Biology, University of Arizona, Tucson, AZ, 85721, USA; Department of Ecology, Behavior, and Evolution, University of California San Diego, San Diego, CA, 92093, USA

**Keywords:** linkage disequilibrium, population genetics, nearly neutral theory, forward time simulation, effective population size, expected heterozygosity

## Abstract

Background selection describes the reduction in neutral diversity caused by selection against deleterious alleles at other loci. It is typically assumed that the purging of deleterious alleles affects linked neutral variants, and indeed simulations typically only treat a genomic window. However, background selection at unlinked loci also depresses neutral diversity. In agreement with previous analytical approximations, in our simulations of a human-like genome with a realistically high genome-wide deleterious mutation rate, the effects of unlinked background selection exceed those of linked background selection. Background selection reduces neutral genetic diversity by a factor that is independent of census population size. Outside of genic regions, the strength of background selection increases with the mean selection coefficient, contradicting the linked theory but in agreement with the unlinked theory. Neutral diversity within genic regions is fairly independent of the strength of selection. Deleterious genetic load among haploid individuals is underdispersed, indicating non-independent evolution of deleterious mutations. Empirical evidence for underdispersion was previously interpreted as evidence for global epistasis, but we recover it from a non-epistatic model.

**SIGNIFICANCE:** As individuals bearing deleterious alleles are removed from a population, other alleles are removed with them, some that are tightly linked near the deleterious allele on a chromosome and some that aren’t linked at all. When the deleterious mutation rate is realistically high, unlinked pairs of loci are a more important influence on the removal of genetic variation. Simulations that assume independent evolution cannot capture removal just by using a lower “effective population size”, because the probabilities of having deleterious alleles on different chromosomes are negatively correlated rather than independent.

## INTRODUCTION

The neutral theory of molecular evolution postulates that i) most genetic diversity observed in natural populations is neutral with respect to an organism’s fitness (Kimura 1968; King and Jukes 1969), and ii) dynamics are well-described by models of a single neutral locus in a population of a specified “effective” population size (Charlesworth 2009; Ewens 2004; Kern and Hahn 2018; Masel 2011). A crucial extension of this theory uses a reduction in the effective population size to incorporate the fact that deleterious mutations depress genetic diversity as they are removed, in a process known as background selection (Charlesworth, Morgan, and Charlesworth 1993). While the removal of a deleterious allele will depress genetic diversity at neutral loci across the entire genome, neutral loci linked to the deleterious allele will be particularly affected (Charlesworth 2012). Indeed, the effect of unlinked background selection is often considered small enough to be ignored (Charlesworth 2012). Here we revisit this assumption given realistically high genome-wide deleterious mutation rates, by using simulations to evaluate previously derived equations for linked vs. unlinked background selection.

In practice, observed genetic diversity at putatively neutral sites is used to estimate the coalescent effective population size *N*_*e*_ as the census size of an idealized population that produces the same neutral genetic diversity given demography (Charlesworth 2009) or background selection (Charlesworth et al. 1993). In an idealized population obeying Wright-Fisher or Moran dynamics, genetic diversity depends only on the product of the neutral mutation rate at that locus and the census size of the population (Kimura 1969). Neutral theory considers mutations that either are strictly neutral or so deleterious that they are purged quickly enough as to leave no impact. Nearly neutral theory retains the binary distinction between rapidly purged versus neutral mutations, but allows the ratio of mutations of these two types to vary among species, according to that species’ value of an effective population size (Ohta 1973). Both models assume many independent single loci.

The problem with this binary distinction and independent loci approach is that slightly deleterious mutations are purged only slowly from populations. During this removal process, they depress genetic variation in the genome (Charlesworth et al. 1993). This depression in genetic variation caused by background selection is typically modeled as a decrease in the effective population size for a neutral locus linked to deleterious variants (Comeron 2014; Hudson and Kaplan 1995; Lohmueller et al. 2011). In a population with no recombination and sufficiently large effect deleterious mutations (*N*_*e*_*s ≫ 1*), the coalescent effective population size would decrease from *N*_*e*_ to *f*_*o*_*N*_*e*_, where *f*_*o*_ is the equilibrium frequency of individuals with no deleterious mutations, because any neutral variants linked to deleterious variants would be doomed (Charlesworth et al. 1993).

Recombination can decouple neutral variants from deleterious variants, reducing the degree to which background selections depresses neutral variation (Cutter and Payseur 2013). For a single neutral locus linked to a single locus where deleterious mutations with fixed heterozygous effect size *sh* occur at rate *u* per diploid individual per generation, and with recombination between the loci occurring at rate *r*, heterozygosity at the neutral locus is reduced by a factor 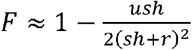 (Hudson and Kaplan 1994).

This result can be straightforwardly extended to any number of deleterious sites linked to the focal neutral site by assuming i) that mutation and recombination rates are uniform across a genomic window centered on the neutral site, ii) that the window is small enough that recombination rate and map length are linearly related, and iii) that there is linkage equilibrium among deleterious variants e.g. because multiple significantly linked deleterious mutations are not present at the same time (Hudson and Kaplan 1995; Nordborg, Charlesworth, and Charlesworth 1996). In this case, in a genomic window we have

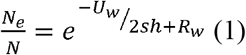

where *U*_*w*_ is the total diploid deleterious mutation rate across the entire window, and *R*_*w*_ is the map length between the ends of the window (Hudson and Kaplan 1995). Assuming a sufficiently large window size (such that *R*_*w*_ *≫ s*), this equation simplifies to

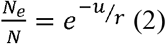

where 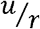 is the ratio of deleterious mutation rate to recombination rate (Charlesworth 2012). With the same assumptions, similar results can be obtained in the case where both deleterious and beneficial mutations occur (Kim and Stephan 2000).

The results above do not apply across different chromosomes or given free recombination between the ends of the window (Hudson and Kaplan 1995). However, deleterious variants do still depress neutral diversity even at unlinked sites: at a bare minimum, a deleterious mutation will eliminate any unique neutral genetic variants in a single individual who dies due to that deleterious mutation. The ratio of *N*_*e*_ to *N* with free recombination has been derived as

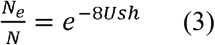

where *U* is the total deleterious mutation rate at unlinked loci and *sh* is the fitness effect of a deleterious allele when in the heterozygous state (Charlesworth 2012).

It has previously been presumed that the reduction in diversity due to unlinked deleterious loci will be much smaller than that from linked loci (Charlesworth 2012). That is, for any given neutral locus, even though *U* across all unlinked deleterious loci is much larger than *U*_*w*_ across the window of linked deleterious loci, 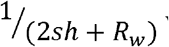 will be so much larger than *8sh* that the latter comparison will overwhelm the former comparison, and linked background selection will dominate. Indeed, models of background selection often simply omit the effects of unlinked background selection (Elyashiv et al. 2016; Ewing and Jensen 2016; McGee et al. 2022; Torres, Szpiech, and Hernandez 2018).

But omitting the effects of unlinked background selection may be a concern given the sheer quantity of deleterious mutations entering populations, and the greater dependence of Equation 3 on this rate. For example, the average number of new deleterious mutations per human is estimated to be at least 2.1 (Lesecque, Keightley, and Eyre-Walker 2012). This estimate is derived from a point mutation rate of 1.1×10^−8^, mutations only being deleterious in the 55% of the 6×10^9^ diploid genome that is not dominated by the remnants of transposable elements, which evolves due to this constraint at 94.3% of the rate. This estimate is conservative because some mutations to transposable element regions are deleterious, because more recent estimates of the human point mutation rate are slightly higher at ∼1.341×10^−8^ (Wang and Obbard 2023), and because non-point mutations and beneficial mutations are neglected. Some therefore argue that deleterious mutation rates are even higher, closer to 10 new deleterious mutations per person in humans (Kondrashov 2017). High deleterious mutation rate estimates are not unique to humans (Haag-Liautard et al. 2007; Popovic et al. 2023).

Here we numerically compare Equations 2 and 3 given human parameters, confirming that unlinked selection has the larger effect. This wouldn’t matter for some purposes, so long as unlinked selection were well described by one-locus models of genetic drift with a lower effective population size. To test this, we perform a multi-locus simulation using the fwdpy11 package (Thornton 2014, 2019), which efficiently handles large numbers of non-neutral mutations in relatively large census size populations (Haller and Messer 2017). These simulations agree reasonably well with past analytic results, confirming the importance of unlinked background selection. More importantly, simulations show that deleterious mutations are not well described by one-locus models, with underdispersion of genetic load among haploid genotypes. This pattern of underdispersion has previously been observed empirically for humans and fruit flies, and interpreted as evidence for epistasis (Sohail et al. 2017); we recover it in a non-epistatic model.

## RESULTS

Unlinked background selection reduces neutral diversity more than linked background selection does (Figure 1A) when the genome-wide deleterious mutation rate *U* is high, specifically when *U > 1*, as is estimated to be the case for humans (Lesecque et al. 2012). This main result is independent of *N* and *s*. A five-fold change in census population size *N* has no significant effect on the 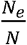 ratio (Figure 1A). This contradicts models that imply that the impact of background selection is likely to be greater in larger populations (Corbett-Detig, Hartl, and Sackton 2015; Cutter and Payseur 2013), although deviations from such models have been previously found (Gillespie 2001; Kaplan, Hudson, and Langle 1989; Santiago and Caballero 2016).

**Figure 1.**
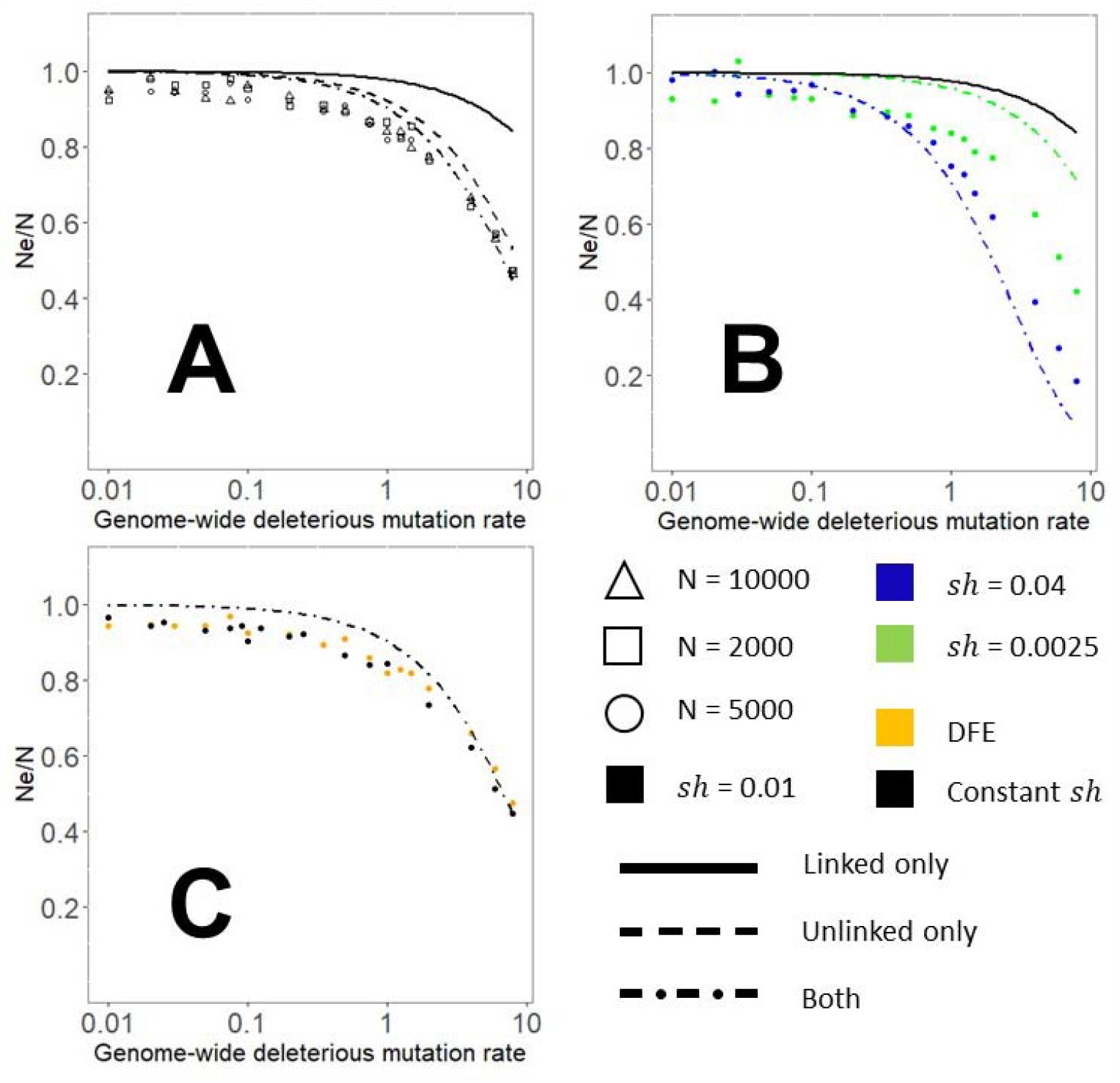
At high deleterious mutation rates, unlinked background selection reduces neutral diversity to a greater extent than linked background selection does. Where not shown, *sh = -0*.*01* and *N* = 5000. Each point represents a single simulation — we chose to allocate computation to a denser grid of parameter values rather than to replicates of the same parameter values. (A) Unlinked background selection alone (dashed line) is a closer match than linked background selection alone (solid line) to the joint model (dot-dashed line) and to simulations (points). Census population size has no effect on the 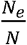 ratio across a five-fold change in simulated census population size. (B) Background selection at high mutation rates is stronger with larger selection coefficients, although our joint model exaggerates this effect. The model with linked background selection alone predicts the opposite dependence on *sh*, and strongly underestimates background selection. (C) The relationship between deleterious mutation rate and *N*_*e*_ is similar whether the effect size of new deleterious mutations is constant versus drawn from a distribution of effect sizes with the same mean value of -0.01.

Larger selection coefficients result in stronger background selection for high *U* but not low *U* (Figure 1B). This is consistent with a dominant role for unlinked background selection (Eq. 3); linked background selection predicts the opposite effect (Eq. 1). Our approximate analytical joint model (see Methods) exaggerates the impact of selection strength; for our focal value of *sh = -0*.*01*, our joint model slightly underestimates background selection (Figure 1A and C), distinctly underestimates it for smaller selection coefficients (Figure 1B green), and slightly overestimates it for large selection coefficients (Figure 1B blue). The underestimation for smaller selection coefficients has been noted before, and likely occurs because the small coefficients are near a weakly deleterious range, violating the assumption that *N*_*e*_*s ≫ 1* (Johri, Charlesworth, and Jensen 2020). A distribution of selective effect sizes behaves similarly to a single *sh* value with the same mean (Figure 1C).

The simulations above assume that deleterious mutations occur uniformly at random across the genome. A more realistic scenario would be for deleterious mutations to be clustered within a functional subset of the genome. We modify our simulations to model genomes where only 10% of the genome is made up of ‘genes’ subject to deleterious mutations. Concentrating deleterious mutations into more tightly linked ‘genes’ results in a smaller reduction in neutral diversity than found in simulations where deleterious mutations occur across the genome (Figure 2, red vs. blue circles). This does not depend on population size (vertical comparison on Figure 2A, B, D).

**Figure 2.**
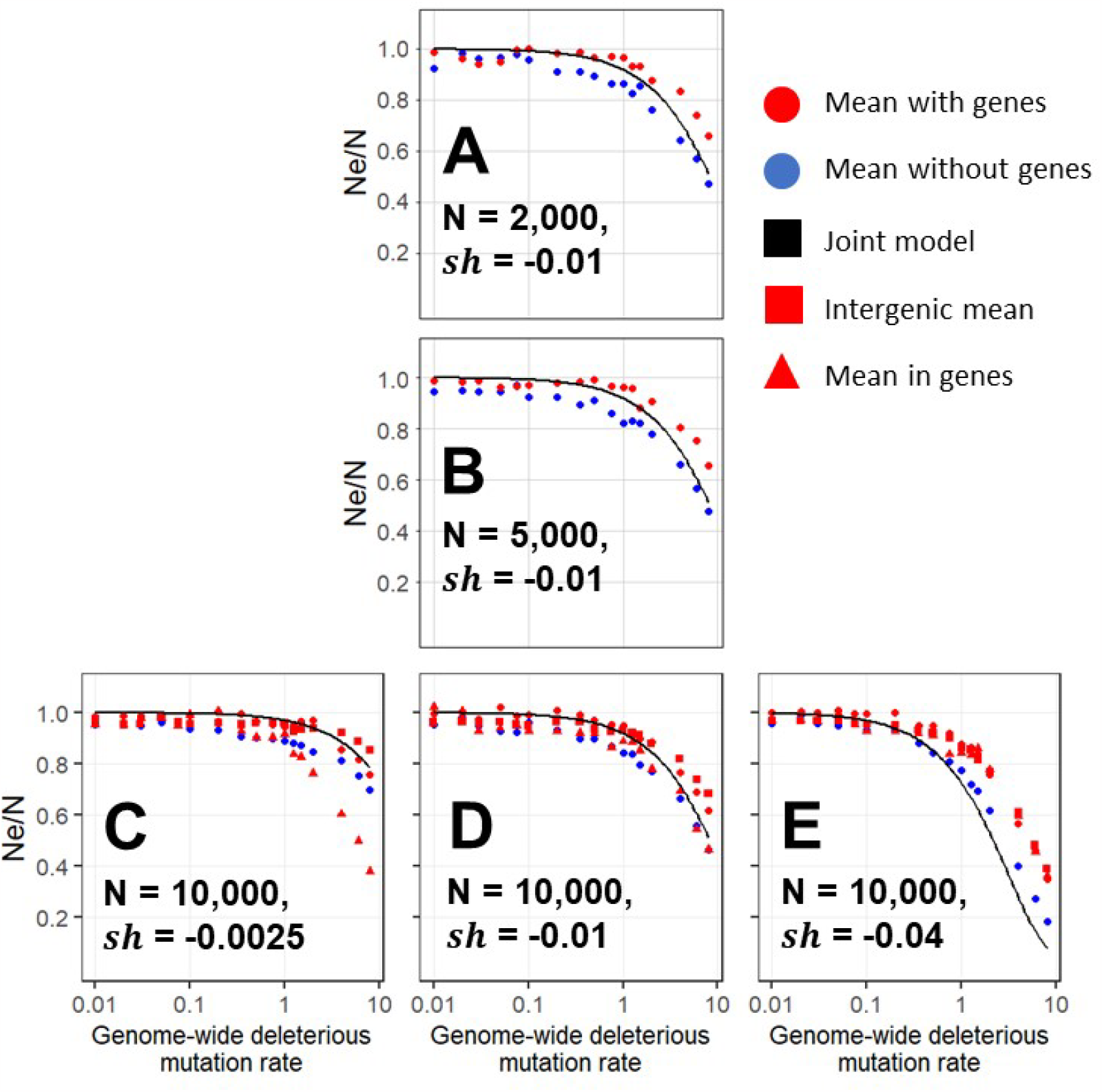
Concentrating deleterious mutations into ‘genes’ slightly weakens overall background selection, and for genic sites, removes the dependence on selection coefficients. As in Figure 1, background selection becomes significant only at high mutation rates, and census population size does not affect results (A vs B vs D). For large selection coefficients, background selection is similar at genic vs intergenic sites (E, red triangles vs squares). For small selection coefficients, the gap is considerable (C, red circles vs squares). The dependence on the strength of selection (circles in C vs D vs E, blue matches Figure 1) does not apply to genic sites (red triangles in C vs D vs E). In all panels, the solid black line is the theoretical expectation incorporating both linked and unlinked background selection, blue dots are simulations where deleterious mutations occur uniformly at random on the genome, and red dots are simulations where deleterious mutations are clustered into ‘genes’ (see Methods). Within the genes condition, neutral sites within genes are indicated as red triangles, and outside them are indicated as red squares.

Simulations quantify the degree to which neutral diversity is more depressed in genes than in intergenic regions (Figure 2, red squares vs. red triangles in panels C, D, and E). At low mutation rates, there is little depression in neutral diversity, with no appreciable difference between linked vs. unlinked sites. There is also little difference with strong selection (Figure 2, panel E); this is expected as we approach the limit at which each deleterious mutation immediately dooms the genome it appears on (Charlesworth et al. 1993). While larger selection coefficients increase overall background selection (discussed above for Fig. 1 and seen for circles in Figure 2 C-E), background selection in genic regions is relatively independent of selection coefficient (red triangles in Figure 2 C-E) and corresponds well to the joint model for *sh = -0*.*01* (Figure 2D).

Strong background selection might be well captured by traditional one-locus models with lower *N*_*e*_, if deleterious sites evolve independently. However, at high deleterious mutation rates, linkage disequilibrium occurs (Barton 1998; Barton and Otto 2005), potentially breaking this assumption. To test for independence, we asked whether the variance in the number of deleterious alleles is lower than that expected from the mean under a Poisson distribution (Figure 3). We find that independent evolution of sites breaks down for *U > 1*. Underdispersion among deleterious alleles has been found empirically for humans and fruit flies (Lee 2022; Sohail et al. 2017) (both with *U > 1* (Haag-Liautard et al. 2007; Lesecque et al. 2012)). The index of dispersion was 0.94 for missense variants in the human “crucial genome” in which these variants are most likely to be deleterious (Sohail et al. 2017). Note that Sohail et al. (2017) and Lee (2022) interpreted underdispersion as evidence for synergistic epistasis, but we find underdispersion even in a non-epistatic model.

**Figure 3.**
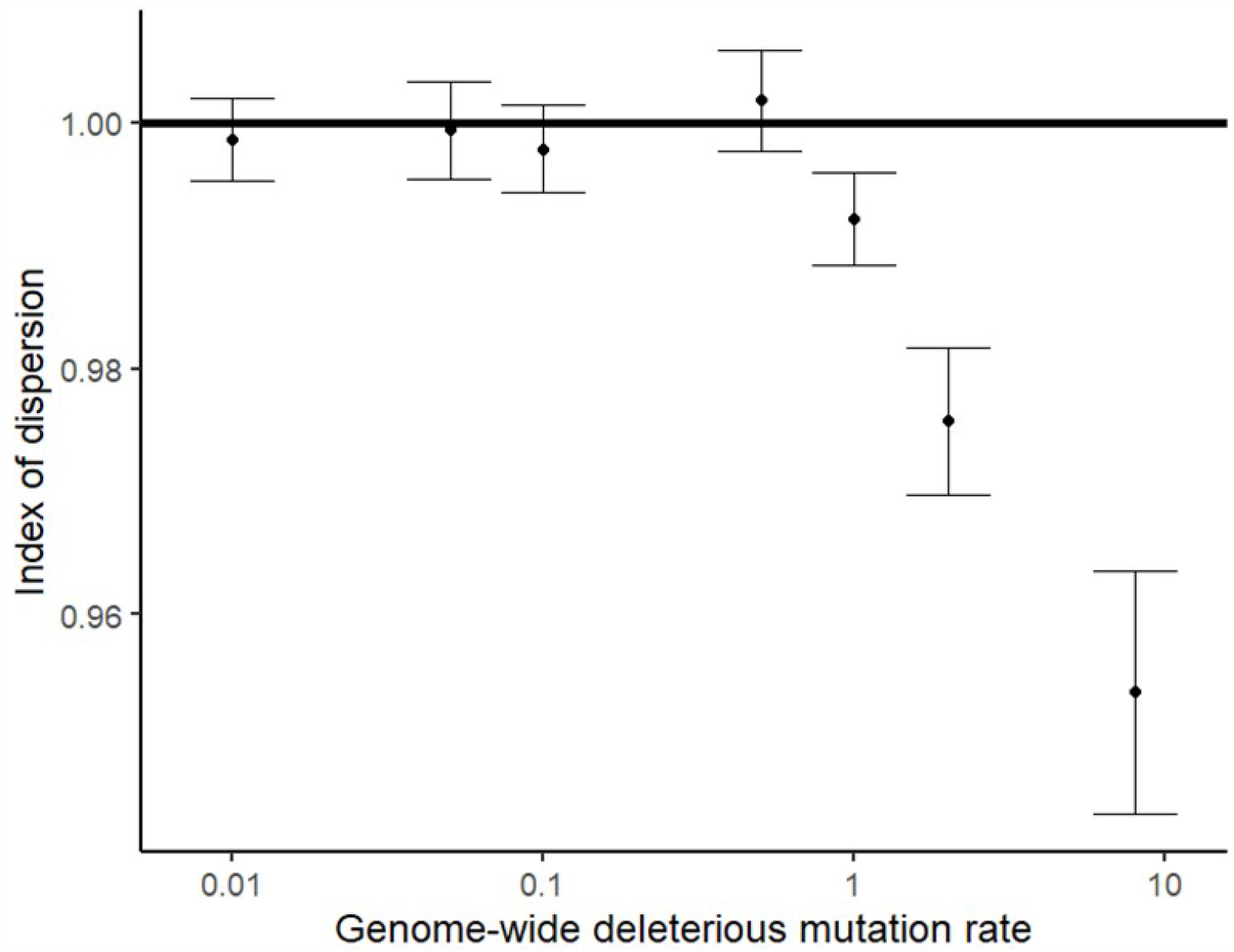
Deleterious mutation rates above 1 create non-independence among segregating sites. The index of dispersion is calculated as the variance in the number of deleterious alleles per haplotype divided by the mean number of deleterious alleles per haplotype. A haplotype is defined as all the chromosomes from one parent, i.e. each individual contains two haplotypes. A Poisson distribution has an index of dispersion of 1, shown as a horizontal line. Points are the mean index of dispersion from eleven replicate simulations with *N* = 10,000, *sh* = −0.01, and bars show the standard error in the estimate of the mean.

## DISCUSSION

High deleterious mutation rates are well established empirically in humans and a variety of other species. Our simulations confirm previous analytic results, showing that when deleterious mutation rates are realistically high, unlinked background selection reduces neutral diversity more than linked background selection does. These findings do not depend on the census population size. Background selection is stronger with larger selection coefficients when deleterious sites are distributed uniformly at random, but when deleterious mutations are clustered into genes, this dependence disappears at neutral sites within genic regions. Existing analytical models exaggerate the dependence on selection coefficients, highlighting our incomplete understanding of unlinked background selection. The original view was that high-*s* mutations at sites that are unlinked to the focal site would exclude an effectively random set of individuals in each generation from the effective population size. This heuristic does not easily explain our findings. However, the approximate fit of our simulation results to models suggest that we are not operating in the interference selection regime characterized by Good *et al*. (2014).

We focused on the index of dispersion as our genome-wide metric of linkage disequilibrium. There have been recent attempts to disentangle pair-wise measures of linkage disequilibrium from their dependence on allele frequency (Garcia and Lohmueller 2021; Good 2022; Potapova and Kondrashov 2023), and the complexities of unphased data (Ragsdale and Gravel 2020). It would be interesting for such work to include unlinked controls in future.

Our multilocus simulations neglect some population features known to affect neutral diversity (e.g. adaptive evolution (Maynard Smith and Haigh 1974) and temporal changes in population size (Torres et al. 2020)), and simplify others (e.g. variation in dominance coefficients among deleterious variants (Gilbert et al. 2020) and heterogeneity in recombination rates (Kulathinal et al. 2008)). The purpose of our simulations is to isolate the effects of background selection with high mutation rates, rather than to accurately reflect the genetics of specific populations. Incorporating additional complications into the model might change the quantitative strength of background selection. However, we do not expect adding new complications to change the broader conclusion that unlinked background selection cannot be safely ignored.

It is already known that the effects of linked background selection are critical for explaining differences in neutral genetic diversity among genomic regions, and that failure to do so is a problem for demographic inference (Ewing and Jensen 2016; Johri et al. 2021; Pouyet et al. 2018). Our results bring up the possibility that unlinked background selection could also be important. At low mutation rates (*U* ≪ 1, see Figure 3), the effects of unlinked background selection are trivial in magnitude. But at realistically high mutation rates (*U* ≫ 1), unlinked background selection will result in non-independent evolution among sites. Non-independence among loci cannot be accounted for with a fudge factor in the effective population size of a one-locus model. In a one-locus model, random changes to allele frequencies are modelled as white noise, i.e. there is no autocorrelation over time. When sites are not independent, being on a bad/good genetic background in one generation will predict being on a bad/good genetic background in the next generation, producing colored noise (Masel 2011). We therefore cannot assume that a one-locus model with lower *N*_*e*_ can accurately capture background selection at unlinked loci.

To save computational cost, previous simulations of background selection treated only a section of a chromosome. The local deleterious mutation rate is set to a value corresponding to high genome-wide *U*, but mutations outside the local window are neglected. We have shown that neglecting them is problematic. This could matter both in the context of demographic inference, and when the goal is to use neutral diversity to distinguish between the effects of linked background selection vs. selective sweeps (Elyashiv et al. 2016; McVicker et al. 2009; Murphy et al. 2022). Neglecting unlinked background selection is also a concern for papers investigating only negative selection; e.g. Torres *et al*. (2018) and Beissinger *et al*. (2016) look for differences in background selection between human or between maize populations, but their measure of background selection assumes a reference class of neutral sites that are unaffected, to be compared to sites subject to strong background selection.

Another common practice in evolutionary simulations is to rescale parameters (e.g. *N, s, U, r*) in a manner that keeps products of interest (e.g. *Ns, Nr*) constant (Campos and Charlesworth 2019; Comeron and Kreitman 2002; Hill and Robertson 1966; Hoggart et al. 2007; Kaiser and Charlesworth 2009; Uricchio and Hernandez 2014). We found that the strength of background selection is independent of *N*, but is stronger for large *s* (at least outside of genic regions). Reducing *N* while increasing *s* will therefore exaggerate the effect of background selection, i.e. the reduction in *N*_*e*_ will be greater than that expected as proportional to the reduction in simulated *N*. The net result of rescaling as it is normally conducted would thus be a reduction in *sN*_*e*_ in populations with high *U*.

Accounting for background selection is often conceived as a necessary step prior to inferring demography and/or adaptation. But unlinked background selection in species with high mutation rates globally suppresses neutral diversity to such a degree that it is worthy of scientific attention in its own right. E.g. discussions of whether background selection is sufficient to resolve Lewontin’s paradox (Buffalo 2021) should consider high *U*_*d*_ and the magnitude of the unlinked background selection that it causes.

To the extent that currently neutral genetic diversity might become relevant in new environments, background selection imposes a genome-wide penalty caused by the constant deluge of deleterious mutations. Massive background selection is thus a sister question to that of overwhelming deleterious mutation load, which has not yet been fully resolved (Agrawal and Whitlock 2012; Goyal et al. 2012; Kondrashov 1995, 2017).

## MATERIALS AND METHODS

### Joint model

To calculate the expected combined effects of background selection at linked and unlinked sites, we simply multiply together the respective reductions in neutral diversity from Equations 2 and 3. To avoid double counting, we subtract from unlinked selection those mutations that fall within a window of presumed linked selection. Considering constant *sh* for convenience, this joint model is given by

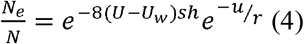

where *U* is the total genome-wide deleterious mutation rate. We assume 20,000 windows (this number is chosen to roughly match the number of genes in humans); using different window numbers has minimal impact (Supplementary Figure 1A). Results are qualitatively the same if we use a joint model constructed from equations 1 and 3, with slightly more dependence on window size (Supplementary Figure 1B).

### Multi-locus simulations

All simulations were written in Python using fwdpy11 (Thornton 2014, 2019). We simulated populations of *N* diploid individuals undergoing selection against deleterious mutations using a standard Wright-Fisher infinite-sites model for 10*N* generations. Each individual’s genome was made up of 23 chromosomes of equal length, with recombination occurring via exactly two cross-overs per chromosome, matching data for humans (Pardo-Manuel De Villena and Sapienza 2001).

Deleterious mutations occur with genome-wide rate *U*. In the “no genes” condition, they are located uniformly at random along the chromosomes, while in the “genes” condition they occur only in “genes”. We simulate 1,000 genes, accounting for 10 percent of the genome, interspersed at regular intervals throughout the genome. Assumptions about “genes” were not chosen to be representative of any particular species, but simply to capture the qualitative consequences of clustering the sites that are subject to deleterious mutations and hence background selection.

A recent study of a large sample of modern European humans estimated a gamma distribution of fitness effects of new non-synonymous mutations with mean *sN*_*e*_ = −224.33 and *2N*_*e*_ = 23,646, implying a mean *sh*≈ -0.01 (Kim, Huber, and Lohmueller 2017). In our main results, we simplify to use a constant *sh* = -0.01 to avoid complications from deleterious mutations with *sN*_*e*_ so near to 1 that they are effectively neutral. We also explore higher and lower values of *s*, and a gamma distribution with the same mean and shape parameter *α* = 0.169 (Kim et al. 2017). All mutations have *h* = 0.5 and fitness is calculated multiplicatively with no epistasis.

While our forward-time simulations track only deleterious mutations, tree-sequence recording (Kelleher et al. 2018) during the simulation allows neutral mutations to be projected backwards onto the genealogical histories of different genomic regions, enabling us to later compute neutral genetic diversity and hence effective population size. In all simulations, neutral mutations occur uniformly at random on the entire genome at an arbitrary rate 10^−4^ per genomic ‘unit’, for a total rate of 0.23 per genome. This low value provides sufficient resolution of *N*_*e*_ at low computational cost. We use msprime (Kelleher, Etheridge, and McVean 2016) to calculate neutral diversity *e* on the resulting tree sequence using an infinite-alleles model, and then calculate the effective population size for a simulation using 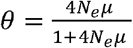 and solving for *N*_*e*_. When calculating effective population size within genes vs. intergenic regions (red squares vs. triangles in Figure 2C-E) we use an infinite-sites model to avoid complications with the distribution of finite neutral sites in genes vs. intergenic regions. In these cases, we calculate the effective population size using *θ* = 4*N*_*e*_*μ* and solving for *N*_*e*_.

We simulated census population sizes *N* ranging from 2,000 to 10,000. This is compatible with the range of inferred estimates for human effective population sizes (McEvoy et al. 2011; Takahata 1993; Tenesa et al. 2007). We calculate the ‘observed’ value of 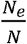 from the neutral diversity, to compare with analytical expectations.

## DATA AVAILABILITY

Simulation code written in python, and graphs produced with R. Scripts available on github at www.github.com/MaselLab/BackgroundSelection.

## ACKNOWLEDGEMENTS

We thank Bruce Walsh for pointing us to Hudson & Kaplan’s analytical results on linked background selection that sparked this work; Denis Roze for a wealth of advice (including pointing us to the equation for unlinked background selection), detailed comments on the manuscript, and catching many errors on our part; two anonymous reviewers of an earlier version of this work who also pointed us to unlinked background selection; Kevin Thornton for pointing out a possible connection with implausibly high empirical estimates of the deleterious mutation rate, and for assistance with fwdpy simulations; and Ryan Gutenkunst, Parul Johri, and David Enard for helpful discussions.

## BIBLIOGRAPHY

Agrawal, Aneil F., and Michael C. Whitlock. 2012. ‘Mutation Load: The Fitness of Individuals in Populations Where Deleterious Alleles Are Abundant’. Annu. Rev. Ecol. Evol. Syst 43:115–35. doi: 10.1146/annurev-ecolsys-110411-160257

Barton, N. H. 1998. ‘The Effect of Hitch-Hiking on Neutral Genealogies’. Genetical Research 72(2):123–33. doi: 10.1017/S0016672398003462

Barton, N. H., and Sarah P. Otto. 2005. ‘Evolution of Recombination Due to Random Drift’. Genetics 169(4):2353–70. doi: 10.1534/genetics.104.032821

Beissinger, Timothy M., Li Wang, Kate Crosby, Arun Durvasula, Matthew B. Hufford, and Jeffrey Ross-Ibarra. 2016. ‘Recent Demography Drives Changes in Linked Selection across the Maize Genome’. Nature Plants 2(7):1–7. doi: 10.1038/NPLANTS.2016.84

Buffalo, Vince. 2021. ‘Quantifying the Relationship between Genetic Diversity and Population Size Suggests Natural Selection Cannot Explain Lewontin’s Paradox’. ELife 10:1–30. doi: 10.7554/eLife.67509

Campos, José Luis, and Brian Charlesworth. 2019. ‘The Effects on Neutral Variability of Recurrent Selective Sweeps and Background Selection’. Genetics 212(1):287–303. doi: 10.1534/genetics.119.301951

Charlesworth, B., M. T. Morgan, and D. Charlesworth. 1993. ‘The Effect of Deleterious Mutations on Neutral Molecular Variation’. Genetics 134(4):1289–1303

Charlesworth, Brian. 2009. ‘Effective Population Size and Patterns of Molecular Evolution and Variation’. Nature Reviews Genetics 10(3):195–205. doi: 10.1038/nrg2526

Charlesworth, Brian. 2012. ‘The Effects of Deleterious Mutations on Evolution at Linked Sites’. Genetics 190(1):5–22. doi: 10.1534/genetics.111.134288

Comeron, Josep M. 2014. ‘Background Selection as Baseline for Nucleotide Variation across the Drosophila Genome’. PLoS Genetics 10(6):e1004434. doi: 10.1371/journal.pgen.1004434

Comeron, Josep M., and Martin Kreitman. 2002. ‘Population, Evolutionary and Genomic Consequences of Interference Selection’. Genetics 161(1):389–410. doi: 10.1093/genetics/161.1.389

Corbett-Detig, Russell B., Daniel L. Hartl, and Timothy B. Sackton. 2015. ‘Natural Selection Constrains Neutral Diversity across A Wide Range of Species’. PLoS Biology 13(4):1–25. doi: 10.1371/journal.pbio.1002112

Cutter, Asher D., and Bret A. Payseur. 2013. ‘Genomic Signatures of Selection at Linked Sites: Unifying the Disparity among Species’. Nature Reviews Genetics 14(4):262–74. doi: 10.1038/nrg3425.Genomic

Elyashiv, Eyal, Shmuel Sattath, Tina T. Hu, Alon Strutsovsky, Graham McVicker, Peter Andolfatto, Graham Coop, and Guy Sella. 2016. ‘A Genomic Map of the Effects of Linked Selection in Drosophila’. PLoS Genetics 12(8):e1006130. doi: 10.1371/journal.pgen.1006130

Ewens, Warren. 2004. Mathematical Population Genetics. 2nd ed. New York: Springer US

Ewing, Gregory B., and Jeffrey D. Jensen. 2016. ‘The Consequences of Not Accounting for Background Selection in Demographic Inference’. Molecular Ecology 25(1):135–41. doi: 10.1111/mec.13390

Garcia, Jesse A., and Kirk E. Lohmueller. 2021. ‘Negative Linkage Disequilibrium between Amino Acid Changing Variants Reveals Interference among Deleterious Mutations in the Human Genome’. PLoS Genetics 17(7):1–25. doi: 10.1371/journal.pgen.1009676

Gilbert, Kimberly J., Fanny Pouyet, Laurent Excoffier, and Stephan Peischl. 2020. ‘Transition from Background Selection to Associative Overdominance Promotes Diversity in Regions of Low Recombination’. Current Biology 30(1):101–107.e3. doi: 10.1016/j.cub.2019.11.063

Gillespie, John H. 2001. ‘Is the Population Size of a Species Relevant To Its Evolution?’ Evolution 55(11):2161. doi: 10.1554/0014-3820(2001)055[2161:itpsoa]2.0.co;2

Good, Benjamin H. 2022. ‘Linkage Disequilibrium between Rare Mutations’. Genetics 220(4). doi: 10.1093/genetics/iyac004

Good, Benjamin H., Aleksandra M. Walczak, Richard A. Neher, and Michael M. Desai. 2014. ‘Genetic Diversity in the Interference Selection Limit’. PLoS Genetics 10(3). doi: 10.1371/journal.pgen.1004222

Goyal, Sidhartha, Daniel J. Balick, Elizabeth R. Jerison, Richard A. Neher, Boris I. Shraiman, and Michael M. Desai. 2012. ‘Dynamic Mutation–Selection Balance as an Evolutionary Attractor’. Genetics 191:1309–19. doi: 10.1534/genetics.112.141291

Haag-Liautard, Cathy, Mark Dorris, Xulio Maside, Steven Macaskill, Daniel L. Halligan, Brian Charlesworth, and Peter D. Keightley. 2007. ‘Direct Estimation of per Nucleotide and Genomic Deleterious Mutation Rates in Drosophila’. Nature 445(7123):82–85. doi: 10.1038/nature05388

Haller, Benjamin C., and Philipp W. Messer. 2017. ‘SLiM 2: Flexible, Interactive Forward Genetic Simulations’. Molecular Biology and Evolution 34(1):230–40. doi: 10.1093/molbev/msw211

Hill, W., and A. Robertson. 1966. ‘The Effect of Linkage on Limits to Artificial Selection.’ Genetical Research 8(3):269–94. doi: 10.1017/S001667230800949X

Hoggart, Clive J., Marc Chadeau-Hyam, Taane G. Clark, Riccardo Lampariello, John C. Whittaker, Maria De Iorio, and David J. Balding. 2007. ‘Sequence-Level Population Simulations over Large Genomic Regions’. Genetics 177(3):1725–31. doi: 10.1534/genetics.106.069088

Hudson, Richard R., and Norman L. Kaplan. 1994. ‘Gene Trees with Background Selection’. Pp. 140–53 in Non-neutral Evolution: Theories and Molecular Data, edited by B. Golding.Boston, MA: Springer US

Hudson, Richard R., and Norman L. Kaplan. 1995. ‘Deleterious Background Selection with Recombination’. Genetics 141(4):1605–17

Johri, Parul, Brian Charlesworth, and Jeffrey D. Jensen. 2020. ‘Toward an Evolutionarily Appropriate Null Model[]’: Genetics 215(May):173–92

Johri, Parul, Kellen Riall, Hannes Becher, Laurent Excoffier, Brian Charlesworth, and Jeffrey D. Jensen. 2021. ‘The Impact of Purifying and Background Selection on the Inference of Population History: Problems and Prospects’. Molecular Biology and Evolution 38(7):2986–3003. doi: 10.1093/molbev/msab050

Kaiser, Vera B., and Brian Charlesworth. 2009. ‘The Effects of Deleterious Mutations on Evolution in Non-Recombining Genomes’. Trends in Genetics 25(1):9–12

Kaplan, Norman L., Richard R. Hudson, and Charles H. Langle. 1989. ‘The “Hitchhiking Effect” Revisited’. Genetics 123(4):887–99

Kelleher, Jerome, Alison M. Etheridge, and Gilean McVean. 2016. ‘Efficient Coalescent Simulation and Genealogical Analysis for Large Sample Sizes’. PLoS Computational Biology 12(5):e1004842. doi: 10.1371/journal.pcbi.1004842

Kelleher, Jerome, Kevin R. Thornton, Jaime Ashander, and Peter L. Ralph. 2018. ‘Efficient Pedigree Recording for Fast Population Genetics Simulation’. PLoS Computational Biology 14(11):e1006581

Kern, Andrew D., and Matthew W. Hahn. 2018. ‘The Neutral Theory in Light of Natural Selection’. Molecular Biology and Evolution 35(6):1366–71. doi: 10.1093/molbev/msy092

Kim, Bernard, Christian Huber, and Kirk Lohmueller. 2017. ‘Inference of the Distribution of Selection Coefficients for New Nonsynonymous Mutations Using Large Samples’. Genetics 206:345–61. doi: 10.1534/genetics.116.197145/-/DC1.1

Kim, Yuseob, and Wolfgang Stephan. 2000. ‘Joint Effects of Genetic Hitchhiking and Background Selection on Neural Variation’. Genetics 155(3):1415–27. doi: 10.1093/genetics/155.3.1415

Kimura, Motoo. 1968. ‘Evolutionary Rate at the Molecular Level’. Nature 217:624–26

Kimura, Motoo. 1969. ‘The Number of Heterozygous Nucleotide Sites Maintained in a Finite Population Due to Steady Flux of Mutations.’ Genetics 61(4):893–903

King, Jack Lester, and Thomas H. Jukes. 1969. ‘Non-Darwinian Evolution’. Science 164(3881):788–98

Kondrashov, Alexey S. 1995. ‘Contamination of the Genome by Very Slightly Deleterious Mutations: Why Have We Not Died 100 Times Over?’ Journal of Theoretical Biology 175(4):583–94. doi: 10.1006/jtbi.1995.0167

Kondrashov, Alexey S. 2017. Crumbling Genome: The Impact of Deleterious Mutations on Humans. Hoboken, NJ: John Wiley & Sons, Inc

Kulathinal, Rob J., Sarah M. Bennett, Courtney L. Fitzpatrick, and Mohamed A. F. Noor. 2008. ‘Fine-Scale Mapping of Recombination Rate in Drosophila Refines Its Correlation to Diversity and Divergence’. Proceedings of the National Academy of Sciences of the United States of America 105(29):10051–56. doi: 10.1073/pnas.0801848105

Lee, Yuh Chwen G. 2022. ‘Synergistic Epistasis of the Deleterious Effects of Transposable Elements’. Genetics 220(2). doi: 10.1093/genetics/iyab211

Lesecque, Yann, Peter D. Keightley, and Adam Eyre-Walker. 2012. ‘A Resolution of the Mutation Load Paradox in Humans’. Genetics 191(4):1321–30. doi: 10.1534/genetics.112.140343

Lohmueller, Kirk E., Anders Albrechtsen, Yingrui Li, Su Yeon Kim, Thorfinn Korneliussen, Geng Tian, Emilia Huerta-sanchez, Alison F. Feder, and Niels Grarup. 2011. ‘Natural Selection Affects Multiple Aspects of Genetic Variation at Putatively Neutral Sites across the Human Genome’. PLoS Genetics 7(10):e1002326. doi: 10.1371/journal.pgen.1002326

Masel, Joanna. 2011. ‘Genetic Drift’. Current Biology 21(20):R837–38. doi: 10.1016/j.cub.2011.08.007

Maynard Smith John, and John Haigh. 1974. ‘The Hitch-Hiking Effect of a Favourable Gene’. Genetics Research 23(1):23–35. doi: 10.1017/S0016672308009579

McEvoy, Brian P., Joseph E. Powell, Michael E. Goddard, and Peter M. Visscher. 2011. ‘Human Population Dispersal “Out of Africa” Estimated from Linkage Disequilibrium and Allele Frequencies of SNPs’. Genome Research 21(6):821–29. doi: 10.1101/gr.119636.110

McGee, Ryan Seamus, Olivia Kosterlitz, Artem Kaznatcheev, Benjamin Kerr, and Carl T. Bergstrom. 2022. ‘The Cost of Information Acquisition by Natural Selection’. Biorxiv 2022.07.02. doi: 10.1101/2022.07.02.498577

McVicker, Graham, David Gordon, Colleen Davis, and Phil Green. 2009. ‘Widespread Genomic Signatures of Natural Selection in Hominid Evolution’. PLoS Genetics 5(5):e1000471. doi: 10.1371/journal.pgen.1000471

Murphy, David, Eyal Elyashiv, Guy Amster, and Guy Sella. 2022. ‘Broad-Scale Variation in Human Genetic Diversity Levels Is Predicted by Purifying Selection on Coding and Non-Coding Elements’. ELife 11. doi: 10.7554/eLife.76065

Nordborg, Magnus, Brian Charlesworth, and Deborah Charlesworth. 1996. ‘The Effect of Recombination on Background Selection’. Genetics Research 67(2):159–74

Ohta, Tomoko. 1973. ‘Slightly Deleterious Mutant Substitutions in Evolution’. Nature 246(5428):96–98

Pardo-Manuel De Villena Fernando, and Carmen Sapienza. 2001. ‘Recombination Is Proportional to the Number of Chromosome Arms in Mammals’. Mammalian Genome 12(4):318–22. doi: 10.1007/s003350020005

Popovic, Iva, Lucie Bergeron, Yves-Marie Bozec, Ann-Marie Waldvogel, Samantha Howitt, Katarina Damjanovic, Frances Patel, Maria G. Cabrera, Gert Worheide, Sven Uthicke, and Cynthia Riginos. 2023. ‘High Germline Mutation Rates but Not Extreme Population Size Outbreaks Influence Genetic Diversity in Crown-of-Thorns Sea Stars’. BioRxiv 2023.06.28. doi: 10.1101/2023.06.28.546961

Potapova, Nadezhda, and Alexey S. Kondrashov. 2023. ‘Positive Association between Alleles at Selectively Neutral Loci’. Biorxiv 1–12. doi: 10.1101/2023.06.14.544962

Pouyet, Fanny, Simon Aeschbacher, Alexandre Thiéry, and Laurent Excoffier. 2018. ‘Background Selection and Biased Gene Conversion Affect More than 95% of the Human Genome and Bias Demographic Inferences’. ELife 7:1–21. doi: 10.7554/eLife.36317

Ragsdale, Aaron P., and Simon Gravel. 2020. ‘Unbiased Estimation of Linkage Disequilibrium from Unphased Data’. Molecular Biology and Evolution 37(3):923–32. doi: 10.1093/molbev/msz265

Santiago, Enrique, and Armando Caballero. 2016. ‘Joint Prediction of the Effective Population Size and the Rate of Fixation of Deleterious Mutations’. Genetics 204(3):1267–79. doi: 10.1534/genetics.116.188250

Sohail, Mashaal, Olga A. Vakhrusheva, Jae Hoon Sul, Sara L. Pulit, Laurent C. Francioli, Leonard H. Van Den Berg, Jan H. Veldink, Paul I. W. De Bakker, Georgii A. Bazykin, Alexey S. Kondrashov, and Shamil R. Sunyaev. 2017. ‘Negative Selection in Humans and Fruit Flies Involves Synergistic Epistasis’. Science 356(6337):539–42. doi: 10.1126/science.aah5238

Takahata, N. 1993. ‘Allelic Genealogy and Human Evolution’. Molecular Biology and Evolution 10(1):2–22. doi: 10.1093/oxfordjournals.molbev.a039995

Tenesa, Albert, Pau Navarro, Ben J. Hayes, David L. Duffy, Geraldine M. Clarke, Mike E. Goddard, Peter M. Visscher, Human Genetics Unit, Western General Hospital, Edinburgh Eh, United Kingdom, and Evolutionary Biology. 2007. ‘Recent Human Effective Population Size Estimated from Linkage Disequilibrium’. Cold Spring Harbor Laboratory Press Human 2:520–26. doi: 10.1101/gr.6023607.8

Thornton, Kevin R. 2014. ‘A C++ Template Library for Efficient Forward-Time Population Genetic Simulation of Large Populations’. Genetics 198(1):157–66. doi: 10.1534/genetics.114.165019

Thornton, Kevin R. 2019. ‘Polygenic Adaptation to an Environmental Shift: Temporal Dynamics of Variation under Gaussian Stabilizing Selection and Additive Effects on a Single Trait’. Genetics 213(December):1513–30. doi: 10.1101/505750

Torres, Raul, Markus G. Stetter, Ryan D. Hernandez, and Jeffrey Ross-Ibarra. 2020. ‘The Temporal Dynamics of Background Selection in Nonequilibrium Populations’. Genetics 214(4):1019–30

Torres, Raul, Zachary A. Szpiech, and Ryan D. Hernandez. 2018. ‘Human Demographic History Has Amplified the Effects of Background Selection across the Genome’. PLoS Genetics 14(6):1–27. doi: 10.1371/journal.pgen.1007387

Uricchio, Lawrence H., and Ryan D. Hernandez. 2014. ‘Robust Forward Simulations of Recurrent Hitchhiking’. Genetics 197(1):221–36. doi: 10.1534/genetics.113.156935

Wang, Yiguan, and Darren J. Obbard. 2023. ‘Experimental Estimates of Germline Mutation Rate in Eukaryotes: A Phylogenetic Meta-Analysis’. Evolution Letters 7(4):216–26. doi: 10.1093/evlett/qrad027

