## Supplementary Figure 1 for "Background selection from unlinked sites causes non-independent evolution of deleterious mutations"

**Supplementary Figure 1.** The window size chosen has little effect on the expected genetic diversity from equation 4. Panel A uses equations 2 and 3 (as do most of our figures), while panel B uses equations 1 and 3. Window size makes a negligible difference for the former (almost perfect superposition), and a small difference for the latter, except for inappropriately large windows (the blue lines, showing only 100 windows for the whole genome). With more windows, the two formalizations are indistinguishable. Open shapes show simulation runs for different population sizes, from Figure 1A.
